# Genetic architecture of the murine serum metabolome reveals carboxyl esterases as master regulators of circulating fatty acid metabolism

**DOI:** 10.64898/2026.06.22.733914

**Authors:** Gregory R. Keele, Travis Nemkov, Ariel M. Hay, Matthew Vincent, Callan O’Connor, Daniel Stephenson, Grier P. Page, James C Zimring, Gary A. Churchill, Angelo D’Alessandro

## Abstract

**Background:** The systemic biochemical diversity of circulating metabolites and lipids reflects the integrated effects of genetic variation and environmental exposure. Metabolite quantitative trait locus (mQTL) studies in humans have established gene-metabolite associations, but genetic contributions can be obscured by sex, diet, age, medication use, and environmental exposures. Genetically diverse model systems offer a powerful complementary strategy to isolate genetic contributions to the biochemical diversity of the circulating metabolome.

**Methodology/Principal Findings:** We applied mass spectrometry profiling to serum samples in 541 mice from the Diversity Outbred (DO) population and identified 1,933 mQTL across 240 metabolites, 561 lipids, 43 oxylipins, and 4,465 MS/MS features. Co-mapping QTL, i.e., QTL hotspots, on chromosomes 8 and 17 implicated carboxyl esterase gene clusters (*Ces1* and *Ces2*) as major regulators of circulating lipid remodeling and demonstrated genetic control of circulating protein/peptide-like features at the major histocompatibility complex and complement *C3* loci. QTL hotspots on chromosomes 9 and 10 revealed previously unknown genetic drivers of lipid and amino acid metabolism. Comparisons with matched red blood cell mQTL revealed widespread compartment-specific genetic control.

**Conclusions/Significance:** Collectively, these findings provide a high-resolution map of the genetic regulation of the circulating metabolome, offering mechanistic insights that complement and extend human metabolic genetics.

## INTRODUCTION

Characterizing the circulating metabolome and its variation across individuals has long been central to clinical chemistry [1], especially for diagnosing inborn metabolic disorders [2, 3]. These disorders arise from genetic variation that impairs enzyme function, resulting in abnormal accumulation of metabolites detectable in blood and urine. Early metabolite profiling approaches enabled the diagnoses of metabolic disorders, such as phenylketonuria, maple syrup urine disease, and organic acidemias [4]. These examples underscore how genetic variation can shape circulating metabolite levels and, ultimately disease susceptibility.

The circulating metabolome and lipidome serve as dynamic intermediates linking genetic variation to complex traits and diseases [1]. Genome-wide association studies (GWAS) have identified numerous metabolic quantitative trait loci (mQTL) that regulate metabolite and lipid levels [5] and have clarified pathways relevant to cardiovascular diseases, diabetes, and neurodegenerative disorders [6, 7]. Notable metabolic pathways that have been elucidated through genetic analysis include branched-chain amino acid metabolism, fatty acid metabolism, and lipid trafficking [8].

Human studies, however, remain constrained by limited genetic diversity and heterogeneity across environmental exposures, collectively known as the exposome [9]. Lifestyle, diet, medications, and other behaviors and exposures strongly modulate the circulating metabolome [9, 10]. This multitude of influences complicates efforts to tease genetic effect apart from the environment. Furthermore, functional validation of human GWAS candidates poses an additional barrier to deriving mechanistic insights.

Mice offer a mature mammalian model system for dissecting the genetic architecture of metabolomic and lipidomic traits [11]. Classical inbred mouse strains have yielded major insights into metabolic regulation, including glucose metabolism, lipid processing, and obesity [12–14]. Nonetheless, their limited genetic diversity limits discovery and translational relevance. Multi-parental model systems [15] retain the experimental control of classical strain while capturing genetic complexity approximating natural human populations. The multi-parental inbred Collaborative Cross (CC) [16] and its outbred sister population, Diversity Outbred (DO) [17] mice, possess extensive genetic diversity derived from eight distinct founder strains [18]: A/J (AJ), C57BL/6J (B6), 129S1/SvlmJ (129S), NOD/ShiLtJ (NOD), NZO/HILtJ (NZO), CAST/EiJ (CAST), PWK/PhJ (PWK), and WSB/EiJ (WSB). The DO population reservoir of segregating genetic variation through continuous outbreeding and thus supports high-resolution mapping of complex traits [11, 17]. Studies have previously leveraged the DO population to map genetic loci affecting lipid metabolism, glucose homeostasis and glycolysis, insulin sensitivity, and aging [14–29].

Despite these advancements, the broader genetic networks governing circulating metabolites and lipids remains incompletely understood. Prior studies have mapped individual QTL but have not fully defined the genetic architecture underlying metabolic variation. Here, we employ the DO population to map genetic determinants of serum metabolites and lipids with high resolution. Using targeted and untargeted metabolomics, lipidomics, and oxylipin profiling, we identify new genetic drivers of circulating biochemical diversity. We leverage existing gene and protein expression datasets from other DO cohorts to define and prioritize candidate genes at QTL hotspots. This work advances our understanding of metabolic regulation and provides a foundation for future mechanistic and translational studies.

## RESULTS

To dissect the genetic architecture of the serum metabolome and lipidome, we profiled 541 Diversity Outbred (DO) mice using mass spectrometry-based metabolomics, lipidomics, and oxylipin analyses (**Figure 1A**). Our integrated dataset provided a comprehensive biochemical survey of mouse serum, capturing a total of 240 metabolites, 561 lipids, 43 oxylipins, and 4,465 unidentified MS/MS features after quality control checks (Methods).

**Figure 1.**
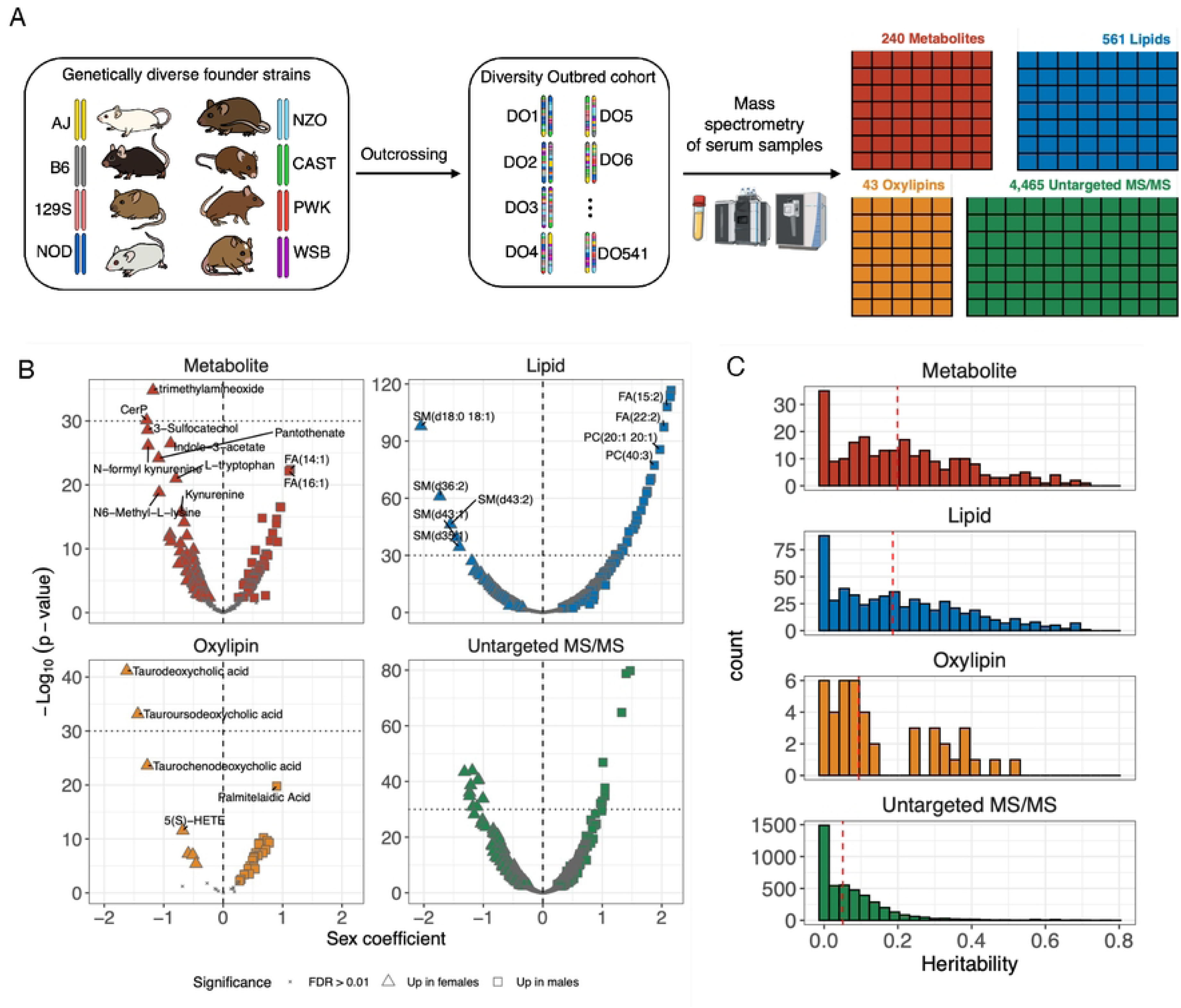
A genetic resource for dissecting the determinants of metabolic and lipidomic variation. (A) Overview of the experimental design, including breakdown of mouse population and serum metabolite profiling. (B) Sex differences as volcano plots for each data layer. Horizontal dotted line at –Log_10_(p-value) = 30 included for reference point across data layers. Vertical dashed line at 0 included for reference. (C) Histograms of narrow sense heritability for each data layer. Red vertical dashed line represents median heritability.

### Sex Differences in Serum Biochemical Features

We first evaluated whether individual biochemical traits differed between the males and females. Significant sex differences (FDR < 0.01) were detected for 119 metabolites, 297 lipids, 30 oxylipins, and 1,238 unidentified MS/MS features (**Figure 1B; Data S1**). Overall, this represented 49.6% of metabolites, 52.9% of lipids, 69.8% of oxylipins, and 27.7% of unidentified MS/MS features. We caution against over-interpreting these findings given that sex was partially confounded with batch (Methods). Nevertheless, consistent patterns emerged between metabolites and lipids: among fatty acids exhibiting sex differences, 14 of 15 metabolites and 26 of 29 lipid species were elevated levels in males.

### QTL Analysis Reveals Distinct Hotspot Loci Driving Serum Biochemical Variation

To assess the cumulative genetic contribution to each trait, we estimated the narrow sense heritability, representing the proportion of variance attributable to additive genetic effects (**Figure 1C**). The median heritability for metabolites and lipids is 19.9% and 18.6%, respectively. Oxylipins and unidentified MS/MS features showed lower heritability comparatively. All data types possessed right-skewed heritability distributions, indicating the presence of highly heritable traits well suited for QTL analysis.

We next performed genome-wide QTL scans across all traits (Methods). A lenient logarithm of odds (LOD) score threshold of 7 was selected to maximize discovery potential and enable detection of genetic loci that regulate many, likely biologically related molecules, i.e., QTL hotspots (**Figure 2A-D**). We detected 163 QTL for metabolites, 324 for lipids, 17 for oxylipins, and 1429 for unidentified MS/MS (**Data S2**). To define QTL hotspots, we used a more stringent LOD score threshold of 10, and then required at least eight QTL to map within a 4 Mbp window of the genome, resulting in 7 QTL hotspots that regulate serum biochemical profiles (**Figure 2E**). For identified molecules, four major hotspots were observed on chromosomes 8 (94 Mbp and 105.5 Mbp), chromosome 9 (96 Mbp), and chromosome 10 (75.5 Mbp). Untargeted MS/MS features also mapped QTL to these hotspots (excluding chromosome 9) and additionally to hotspots on chromosome 1 (171 Mbp) and chromosome 17 (36 Mbp and 58 Mbp).

**Figure 2.**
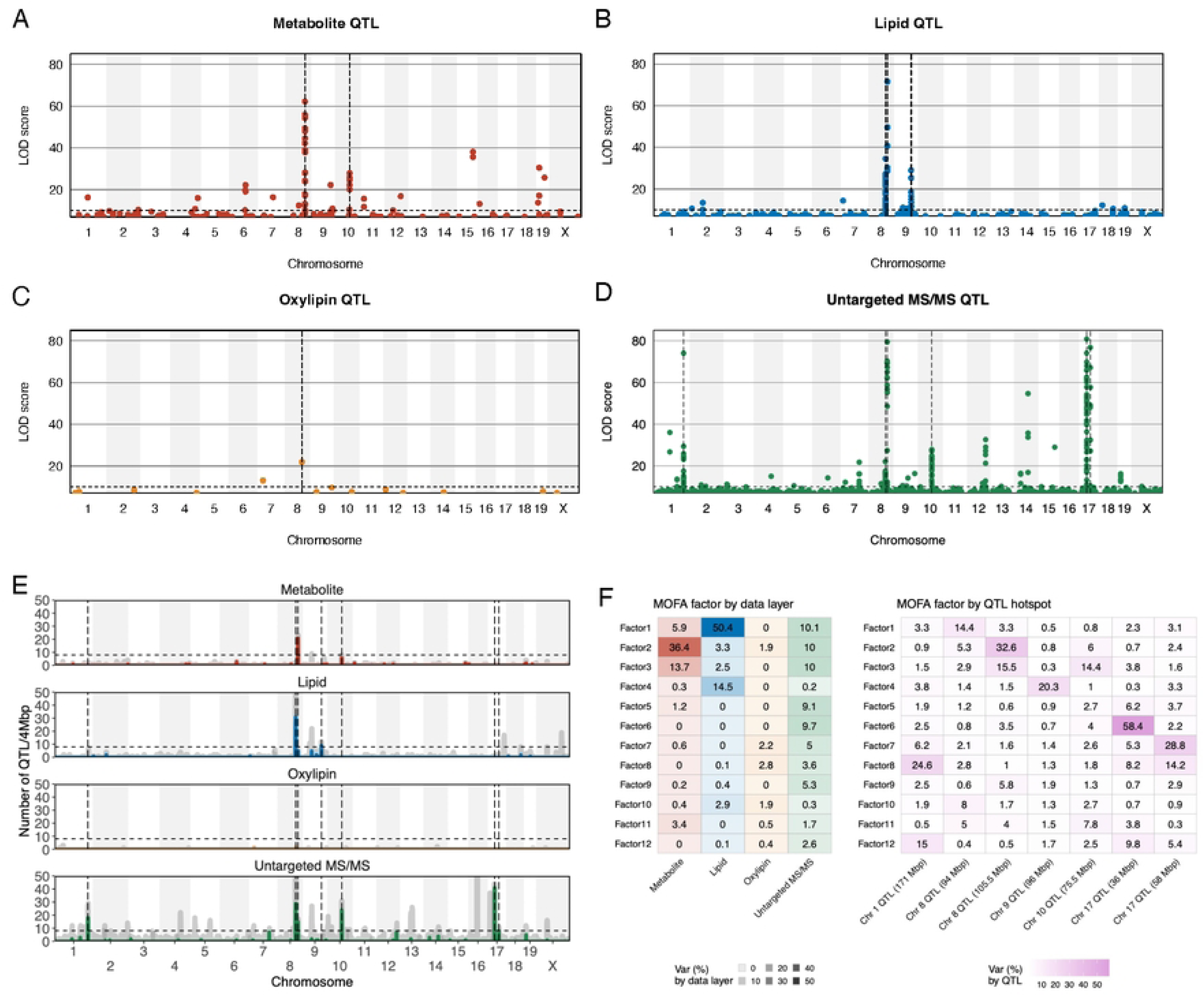
QTL analysis reveals shared QTL hotspots. Manhattan plots of detected QTL (LOD score > 7) for (A) metabolites, (B) lipids, (C) oxylipins, and (D) untargeted MS/MS. Vertical dashed lines represent the seven QTL hotspots mapped in each data layer. Horizontal dashed line represents a LOD score threshold of 10, used to define QTL hotspots. (E) The QTL density (LOD score > 10) across the genome for each data layer. Dark gray density lines represent QTL density with a LOD score threshold of 7. Vertical dashed lines represent the seven QTL hotspots. Horizontal dashed line represents a LOD score threshold of 10, used to define QTL hotspots. (E) MOFA latent factors as heatmaps representing variation partitioned by data layer (left) and by QTL hotspot locus (right).

### MOFA Integration Confirms QTL Hotspots Are Shared Across Serum Biochemical Features

To integrate across omics layers, we performed Multi-Omics Factor Analysis (MOFA) [30] to traits mapping QTL to the seven hotspots. MOFA identified 12 latent factors that each explain >1% of overall variation across the data (**Figure 2F**). For example, latent factor 1 captured significant levels of variation for lipids (50.4%), untargeted MS/MS features (10.1%), and metabolites (5.9%). This factor maps a strong QTL to the chromosome 8 hotspot at 94 Mbp that explains 14.4% of its variation. This is consistent with QTL for metabolites, lipids, and untargeted MS/MS features mapping to the hotspot (**Figure 2A,B,D**). Other factors were highly specific to omics layer, such as latent factor 6, which accounted for 9.7% of the variation for untargeted MS/MS features but 0% for all other layers. This factor captured signal from features that map QTL to the chromosome 17 at 36 Mbp, explaining 58.4% of its variation.

### Chromosome 8 QTL Hotspots: Carboxylesterase Gene Clusters Regulate Circulating Lipid and Carnitine Levels

The chromosome 8 hotspots at 94 Mbp and 105.5 Mbp represent 61 QTL (127 with LOD score > 7) and 41 QTL (45 at LOD score > 7), respectively (**Figure 3A,B**). The hotspot at 94 Mbp, which spans the *carboxylesterase 1* (*Ces1*) gene cluster, mapped QTL for 31 lipids (including lysophosphatidylcholines, lysophosphatidic acids, and phospholipids), one oxylipin (tauroursodeoxycholic acid [TUDCA]), and 29 untargeted MS/MS features. The downstream hotspot at 105.5 Mbp covers the *carboxylesterase 2* (*Ces2*) gene cluster and mapped QTL for 21 metabolites (acyl-carnitines), five lipids (acyl-CoAs), and 15 untargeted MS/MS features. These results indicate that these gene clusters play a central role in lipid hydrolysis and remodeling.

**Figure 3.**
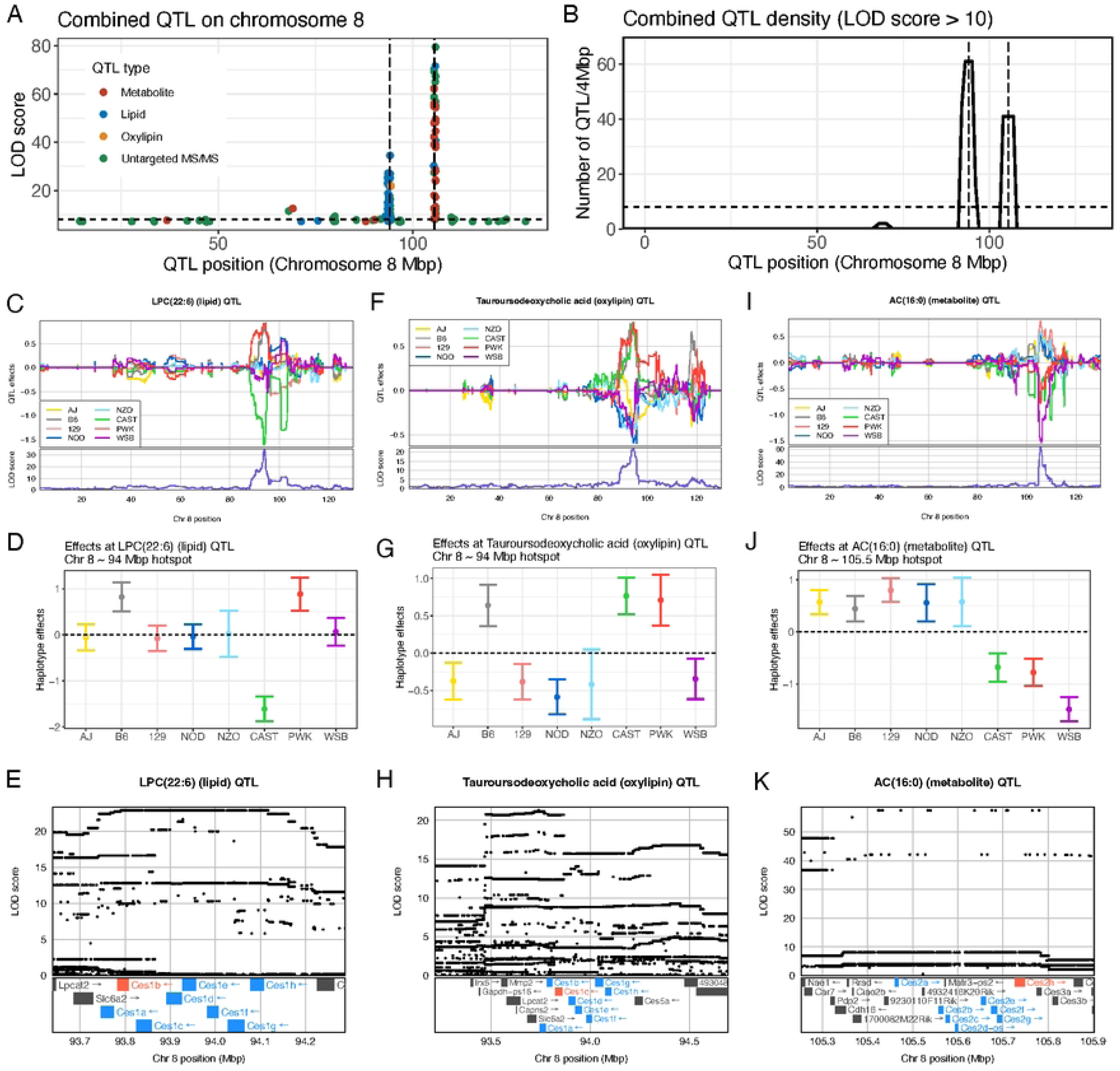
QTL hotspots on chromosome 8 reveal *Ces1* and *Ces2* as significant drivers of serum metabolite and lipid levels. (A) Manhattan plot of detected QTL (LOD score > 7) on chromosome 8. Vertical dashed lines indicate the two QTL hotspots. Horizontal dashed line represents a LOD score threshold of 10, used to define QTL hotspots. (B) The QTL density (LOD score > 10) across chromosome 8 combining data layers. Vertical dashed lines represent the two QTL hotspots. Horizontal dashed line represents a LOD score threshold of 10, used to define QTL hotspots. Aligned haplotype effects and genome scan on chromosome 8 for (C) LPC(22:6), (D) tauroursodeoxycholic acid (TUDCA), and (E) AC(16:0). Haplotype effects with standard error bars at peak QTL locus for (F) LPC(22:6), (G) TUDCA, and (H) AC(16:0). (G) SNP association and genes within the QTL 1.5 LOD drop support interval for (I) LPC(22:6), (J) TUDCA, and (K) AC(16:0). Ces1 and Ces2 family genes of interest highlighted in blue. Top candidates based on local eQTL and pQTL integration highlighted in red. Annotations for gene models were removed for clarity.

The two hotspots exhibited distinct haplotype effects, consistent with different underlying causal variants at each locus (**Figure 3C-H**). The haplotype effects at the *Ces1* locus appear broadly tri-allelic, with B6 and PWK as high alleles, CAST as a low allele, and the other founder strain alleles intermediate (**Figure 3C,D**). The haplotype effects were highly consistent across the QTL mapping to this hotspot, except for TUDCA (**Figure S1A**). The QTL at the *Ces1* locus for TUDCA, a bile acid-derived oxylipin, showed distinct effects that were more consistent with a bi-allelic pattern—B6, CAST, and PWK as high alleles and the founders as low alleles (**Figure 3F,G**). This suggests that distinct genetic variants within the *Ces1* cluster influence lipid and oxylipin profiles. Prioritizing candidate genes and causal variants within a QTL is often challenging due to linkage disequilibrium (LD). For example, *Ces1a*, *Ces1b*, and *Ces1c* each contain hundreds of genetic variants that distinguish B6, CAST, and PWK from the other founder strains (consistent with the TUDCA QTL; **Figure S2J**), including five missense variants in *Ces1b* and seven in *Ces1c*. Disentangling causal variants for the remaining QTL at *Ces1* is even more challenging due to the potentially tri-allelic effects, which imply the involvement of multiple genetic variants. We also note that formally defining how the QTL haplotype effects resolve into functional alleles, i.e., the allelic series, can be assessed through methods like merge analysis [31] and TIMBR [32–34], though these approaches are still subject to assumptions and multiple sources of uncertainty (e.g.,).

Another approach to identifying and prioritizing candidate genes at a QTL is to incorporate gene expression QTL (eQTL) data, i.e., transcriptome-wide association studies (TWAS) [35]. A limitation is that eQTL (or protein expression QTL [pQTL]) are specific to a tissue type, which may not be relevant to the focal QTL’s phenotype, in our case, metabolite, lipid, and oxylipin levels in serum. We surveyed publicly available gene and protein expression datasets from other DO cohorts and evaluated whether local (i.e., *cis*) eQTL and pQTL co-mapped and possessed consistent haplotype effects with the serum QTL hotspots (Methods; **Data S3**). For QTL mapping to the *Ces1* cluster with roughly a tri-allelic effects pattern, *Ces1b* was the strongest candidate gene, possessing local QTL in liver tissue from two DO cohorts with inverted haplotype effects (**Figure S2A,B**). This implies that higher *Ces1b* expression in the liver is associated with lower lipid levels in serum. For the TUDCA QTL, the strongest candidate gene was *Ces1c*, which also displays inverted eQTL effects compared to the serum QTL (**Figure S2D,E**). Notably, the liver local pQTL does not display inverted effects. This is potentially due to the presence of a *Ces1c* coding variant (rs47490976), for which the reference allele is shared by B6, CAST, PWK, and WSB. Coding variants can bias protein abundance quantitation, at times producing artifactual local pQTL that flag the presence or absence of the reference allele. [36]

Contrasting the *Ces1* hotspot, the haplotype effects at the *Ces2* hotspot appear approximately bi-allelic, characterized by high alleles for the wild-derived founder strains (CAST, PWK, and WSB) compared to the classical laboratory strains (**Figure 3J,K**). Unlike the *Ces1* hotspot, all the QTL mapping to the *Ces2* locus have highly correlated haplotype effects (**Figure S1B**). Based on the local eQTL and pQTL in the liver from two other DO cohorts, *Ces2h* is the best candidate (**Figure S2G,H**). Similar to *Ces1b* and *Ces1c* local eQTL, the *Ces2h* local QTL effects are inverted in reference to the serum QTL, suggesting that high *Ces2h* expression in the liver is associated with low serum levels of lipids and carnitines.

We searched the liver lipid and plasma lipid QTL results from a separate DO cohort [37] that has been heavily phenotyped for obesity- and diabetes-relevant traits [11, 38]. The *Ces2* locus mapped two and three QTL for the liver lipids and plasma lipids, respectively. The traits mapping the QTL were four unidentified lipid features and one acyl carnitine (**Figure S1C,D**). The QTL haplotypes effects were highly consistent with our serum results, and the peak association covers the *Ces2* cluster (**Figure S1E,F**). Taken together, the *Ces1* and *Ces2* hotspots demonstrate that genetic perturbation of esterase activity can significantly alter circulating lipid levels.

### Chromosome 9 QTL Hotspot: Identification of a Novel Locus Regulating Lipid Metabolism

The chromosome 9 hotspot at 96 Mbp represented nine QTL (12 with LOD score > 7) (**Figure 4A,B**). This hotspot, which includes the *Gk5* gene, mapped QTL exclusively for lipid traits, including phosphatidylcholines and triglycerides. The haplotype effects followed a consistent (**Figure 4C**) approximately bi-allelic pattern, with NZO, CAST, and PWK carrying alleles associated with lower lipid levels, e.g., PE (O-18:0; 22:6) (**Figure 4D,E**). The region contains many variants that distinguish NZO, CAST, and PWK from the other founder strains; for example, *Gk5* has over 100 such variants. *Gk5* is an intriguing functional candidate because it encodes glycerol kinase 5 and has been minimally characterized in systemic metabolism, but its regulatory influence across several lipid species suggests a potential role in lipid mobilization or synthesis. Notably, *Gk5* is primarily expressed in the skin (sebaceous gland). [39] Identification of this locus could provide a foundation for future functional studies investigating lipid metabolic networks influenced by *Gk5*. Other genes in the region include *Pls1*, *Atr*, *Xrn1*, *Tfdp2*, and *Atp1b3* (**Figure 4F**). Integration of local eQTL and pQTL from a wide range of tissues (**Data S3**) provide some support for *Pls1*, *Atr*, *Gm16794*, *Gm28085*, *Gm38375*, *Gk5*, and *Tfdp2* as candidate genes (**Figure S3A**).

**Figure 4.**
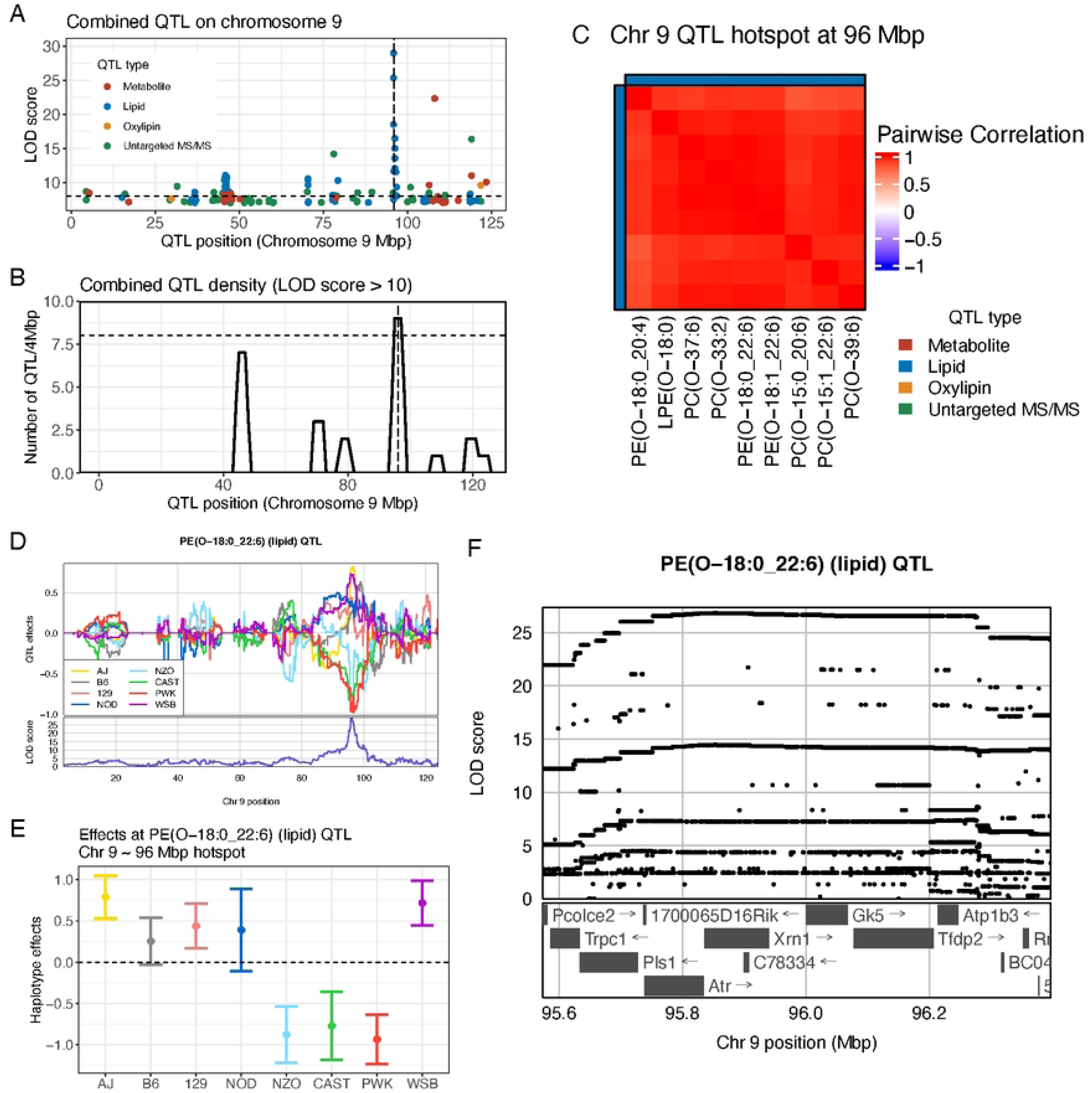
QTL hotspot on chromosome 9 represents novel genetic driver of serum lipid levels. (A) Manhattan plot of detected QTL (LOD score > 7) on chromosome 9. Vertical dashed lines indicate the QTL hotspot. Horizontal dashed line represents a LOD score threshold of 10, used to define QTL hotspots. (B) The QTL density (LOD score > 10) across chromosome 9 combining data layers. The vertical dashed line represents the QTL hotspot. The horizontal dashed line represents a LOD score threshold of 10, used to define QTL hotspots. (C) The heatmap of the correlation matrix of haplotype effects for the QTL mapping to the hotspot. For PE(O-18:0_22:6), (D) aligned haplotype effects and genome scan on chromosome 9, (E) haplotype effects with standard error bars at peak QTL, and (F) SNP association and genes within the QTL 1.5 LOD drop support interval. Annotations for gene models were removed for clarity.

### Chromosome 10 QTL Hotspot: Enrichment for Amino Acid Metabolite QTL

The chromosome 10 hotspot at 75.5 Mbp represents 30 QTL (39 with LOD score > 7) (**Figure 5A,B**). The hotspot mapped QTL for metabolites (including choline, biotin, glycerophosphoethanolamine, and cytidine) and untargeted MS/MS features. The haplotype effects at the locus are similar to *Ces2* hotspot, characterized by wild-derived founder strains (CAST, PWK, and WSB) being distinct from the classical laboratory strains (**Figure 5D,E**). This region contained candidate genes involved in mitochondrial amino acid catabolism and nitrogen homeostasis, including several gamma-glutamyltransferases (*Ggt*), *Ddt*, and *Susd2*. Local eQTL and pQTL integration (**Data S3**) provides some support for *Ggt5*, *Susd2*, *Gm20441*, *Gstt3*, and *Gstt1* (**Figure S3C**).The specificity of metabolites mapping to this locus, together with consistent founder haplotype effects, supports a role for this locus in regulating amino acid utilization pathways. These findings may have relevance for understanding how genetic variation influences nitrogen balance and amino acid turnover, particularly under metabolic stress or dietary perturbation.

**Figure 5.**
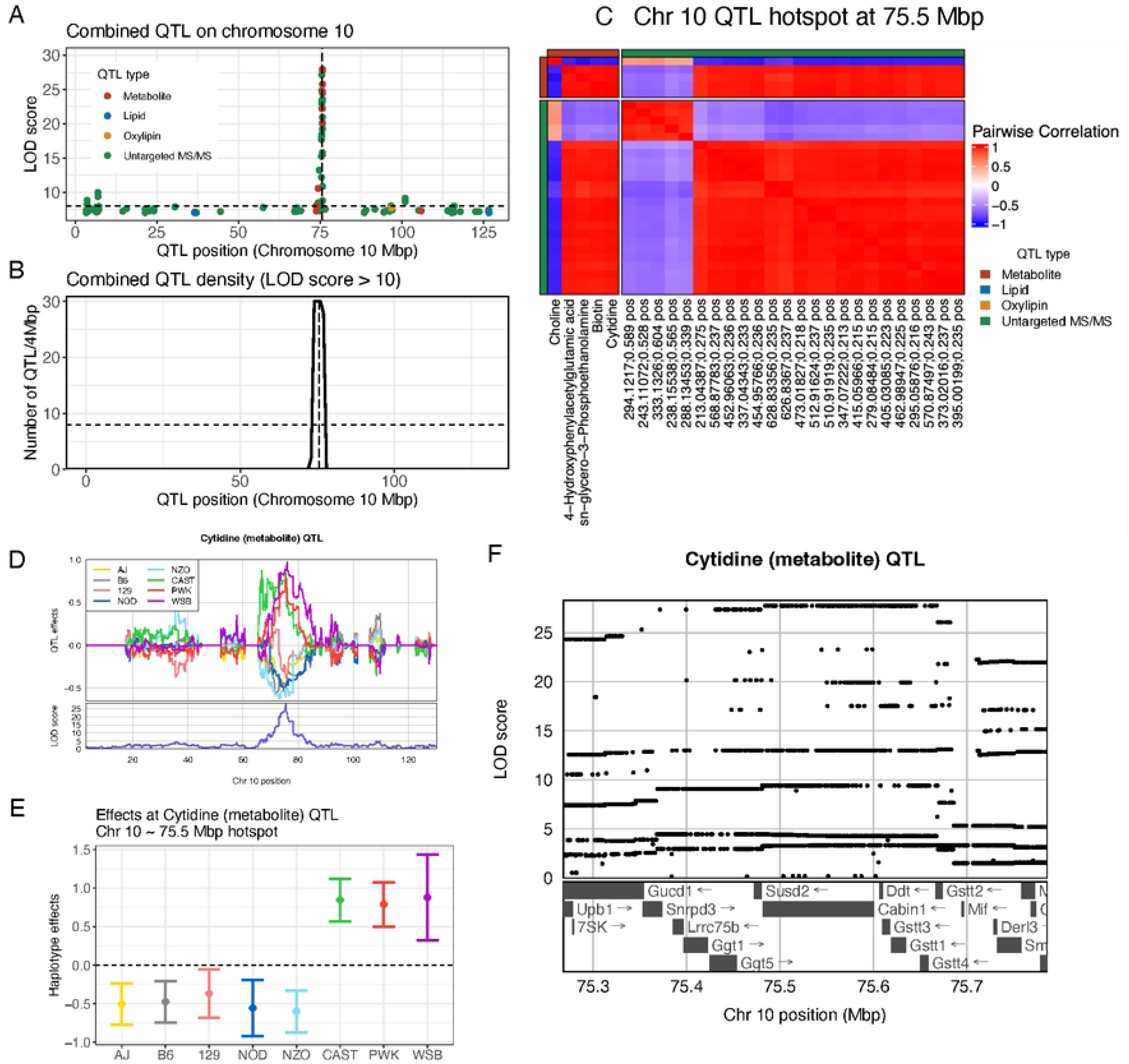
QTL hotspot on chromosome 10 reveals a regulatory hub for amino acid metabolites. (A) Manhattan plot of detected QTL (LOD score > 7) on chromosome 10. Vertical dashed lines indicate the QTL hotspot. Horizontal dashed line represents a LOD score threshold of 10, used to define QTL hotspots. (B) The QTL density (LOD score > 10) across chromosome 10 combining data layers. The vertical dashed line represents the QTL hotspot. The horizontal dashed line represents a LOD score threshold of 10, used to define QTL hotspots. (C) The heatmap of the correlation matrix of haplotype effects for the QTL mapping to the hotspot. For cytidine, (D) aligned haplotype effects and genome scan on chromosome 10, (E) haplotype effects with standard error bars at peak QTL, and (F) SNP association and genes within the QTL 1.5 LOD drop support interval. Annotations for gene models were removed for clarity.

### QTL Hotspots of Untargeted MS/MS Features Reveal Polymorphic Peptide and Proteins Signatures

The untargeted MS/MS features mapped QTL to hotspots shared with identified metabolites and lipids (excluding chromosome 9). They also mapped unique QTL hotspots on chromosome 1 (171 Mbp) and chromosome 17 (36 Mbp and 58 Mbp). The hotspots specific to untargeted MS/MS features were enriched for peptides based on their intact mass ionization state ( [M+2H]+2 and [M+3H]+3), unlike the hotspots that overlap with those for metabolites and lipids (**Figure 6A**). To further evaluate whether these features represented peptide/protein-derived signals, we searched the MS/MS spectra using Mascot and SEQUEST. This analysis yielded 70 high-confidence peptide-spectrum matches, corresponding to 30 peptide entries mapping to 28 unique mouse protein accessions (**Data S4**). Thus, while many untargeted MS/MS features remain unmatched or ambiguous, the identification results support the interpretation that a subset of the QTL hotspots reflects genetically regulated circulating peptide/protein-derived features.

**Figure 6.**
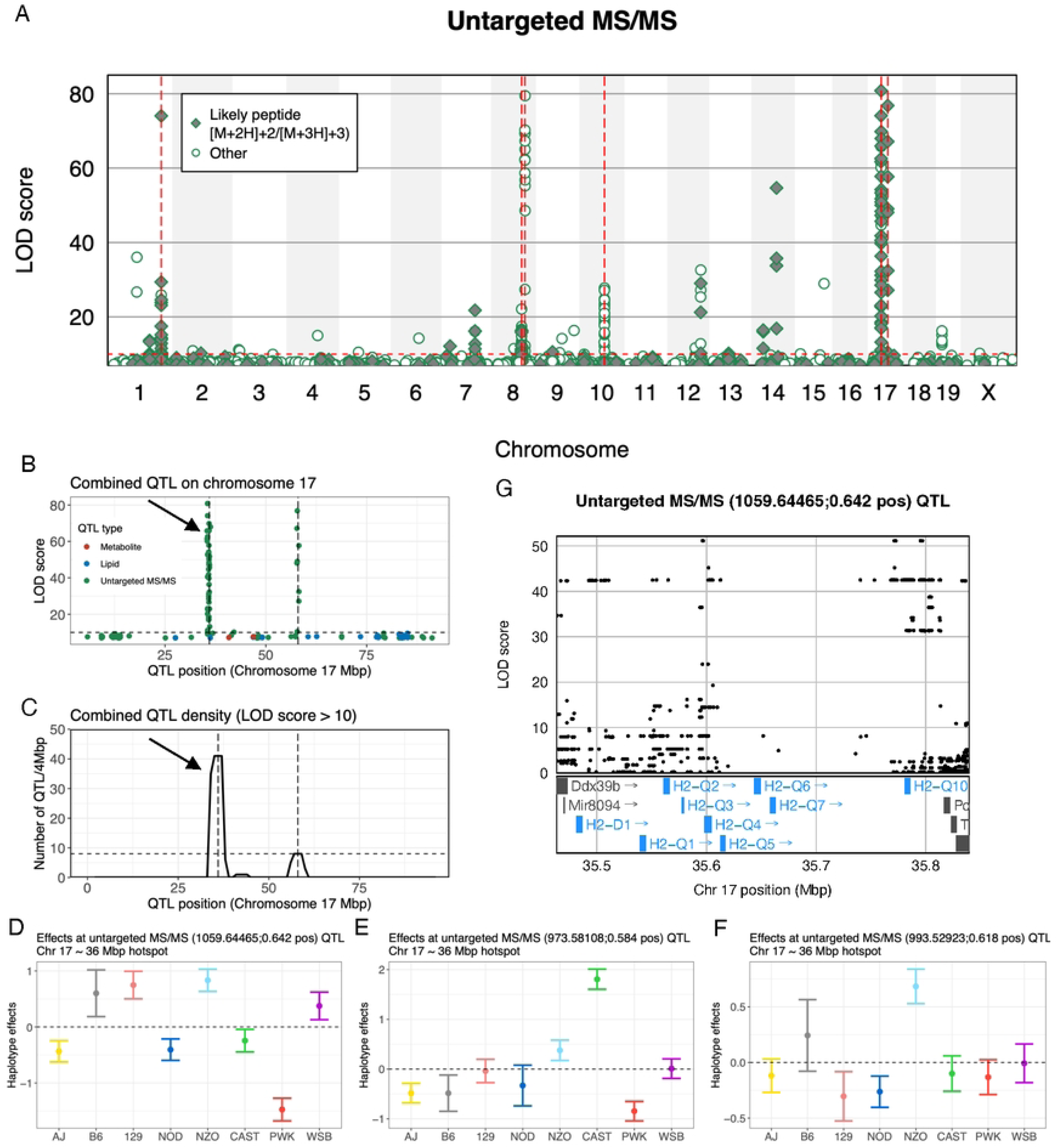
QTL hotspot on chromosome 17 flag high levels of peptide coding variation at the major histocompatibility complex (MHC). (A) Manhattan plot of detected untargeted MS/MS QTL (LOD score > 7). Red vertical dashed lines represent the six QTL hotspots mapped in untargeted MS/MS. Red horizontal dashed line represents a LOD score threshold of 10, used to define QTL hotspots. (B) Manhattan plot of detected QTL (LOD score > 7) on chromosome 17. Vertical dashed lines indicate the QTL hotspots. Horizontal dashed line represents a LOD score threshold of 10, used to define QTL hotspots. Black arrow highlights QTL hotspot at 36 Mbp. (C) The QTL density (LOD score > 10) across chromosome 17 combining data layers. The vertical dashed lines represent the QTL hotspots. The horizontal dashed line represents a LOD score threshold of 10, used to define QTL hotspots. Black arrow highlights QTL hotspot at 36 Mbp. Haplotype effects with standard error bars at peak QTL locus for (D) 1059.64465;0.642 pos, (E) 973.58108;0.584 pos, and (F) 993.52923;0.618 pos. SNP association and genes within the QTL 1.5 LOD drop support interval for 1059.64465;0.642 pos. Candidate genes of interest highlighted in blue. Annotations for gene models were removed for clarity.

The chromosome 1 hotspot at 171 Mbp is composed of QTL for 18 features (28 with LOD score > 7) (**Figure S4A,B**). The haplotype effects at the hotspot are heterogeneous compared to the metabolite-and lipid-centric hotspots (**Figure S4C**), consistent with the possibility that several QTL reflect segregating coding variation in serum peptide/protein-derived species. This is likely because the QTL represent segregating coding variation [40] in peptides in the DO population. For example, the QTL for 1043.42348;0.614_pos displayed an approximately bi-allelic haplotype effects pattern with B6, 129S, NOD, and CAST in a high group and AJ, NZO, PWK, and WSB in a low group (**Figure S4D,E**) compared to 1582.35198;0.647_pos, which also appears largely bi-allelic, but with WSB as a high group compared to the other strains (**Figure S4G,H**). These patterns match the strain distribution patterns for missense variants in *Apoa2*, which is found within the QTL region: rs8258232 (B6, 129S, NOD, CAST vs AJ, NZO, PWK, WSB) and rs242635338 (AJ, B6, 129S, NOD, NZO, CAST, PWK vs WSB). The likely identity of these MS/MS features are polymorphic peptides carrying these specific coding variants.

### Chromosome 17: Coding Variation Signature at Major Histocompatibility Complex (MHC)

One of the largest QTL hotspots was observed on chromosome 17 at 36 Mbp, with 14 untargeted MS/MS mapping QTL (44 with LOD score > 7) to a broad region between 32.3 – 35.5 Mbp (**Figure 6B,C**), which spans the MHC class II locus. Like the chromosome 1 hotspot, the untargeted MS/MS features mapping QTL to the MHC locus likely represent peptide fragments, potentially originating from antigen presentation pathways or proteolytic processing. This interpretation is supported by the more complex multi-allelic haplotype effects observed in the region (**Figure 6D-F**; **Figure S5**), where multiple founder strains (more than two) exhibit distinct alleles, in contrast to the simpler bi-allelic patterns observed for candidate missense mutations at the chromosome 1 hotspot. This complexity, a reflection of a highly genetically heterogenous region that is poorly captured in reference genomes, makes it impossible to pinpoint the specific genetic variants underlying the QTL effects. Nevertheless, the presence of these QTL suggests an interface between immune surveillance and circulating peptide profiles.

A second QTL hotspot for untargeted features was identified further downstream on chromosome 17 at 58 Mbp, consisting of eight untargeted MS/MS QTL (11 with LOD score > 7) (**Figure S6B,C**). Of the eight features that comprise the hotspot, seven map strong QTL (LOD score > 20) and were peptide-like based on ionization state. Notably, the region includes the *C3* complement gene. The haplotype effects are highly bi-allelic, distinguishing the wild-derived founder strains (CAST, PWK, WSB) from the classical laboratory strains (**Figure S6D,E**), consistent with being a signature of coding variation. Examination of missense variants in the region identified rs107650602 and rs52547087 as candidate coding variants in *C3*. Together, these results suggest genetic variation in immune genes may influence serum peptide abundance through mechanisms such as complement activation or proteolytic degradation.

### Serum-to-RBC Metabolite Comparisons Identify Limited Conserved Genetic Regulation

To assess the extent to which our QTL were specific to circulating serum or shared with an intracellular environment, we compared our serum results with our previously published red blood cell (RBC) QTL from the same mouse cohort [20]. We note that the overlap in mice is not complete (477 mice in common for metabolites and lipids and 303 for oxylipins), though comparisons were made using summary statistics and therefore include all the data. Comparison of metabolite QTL revealed that the serum QTL were largely distinct from the RBC QTL (**Figure 7A,B**). In particular, the carnitine-enriched hotspot on chromosome 8 at the *Ces2* locus in serum differs from the carnitine-enriched hotspot on chromosome 11 at the *Slc22a5* locus in RBCs [22]. No QTL hotspots overlapped between serum and RBCs, and no lipid or oxylipin QTL were shared.

**Figure 7.**
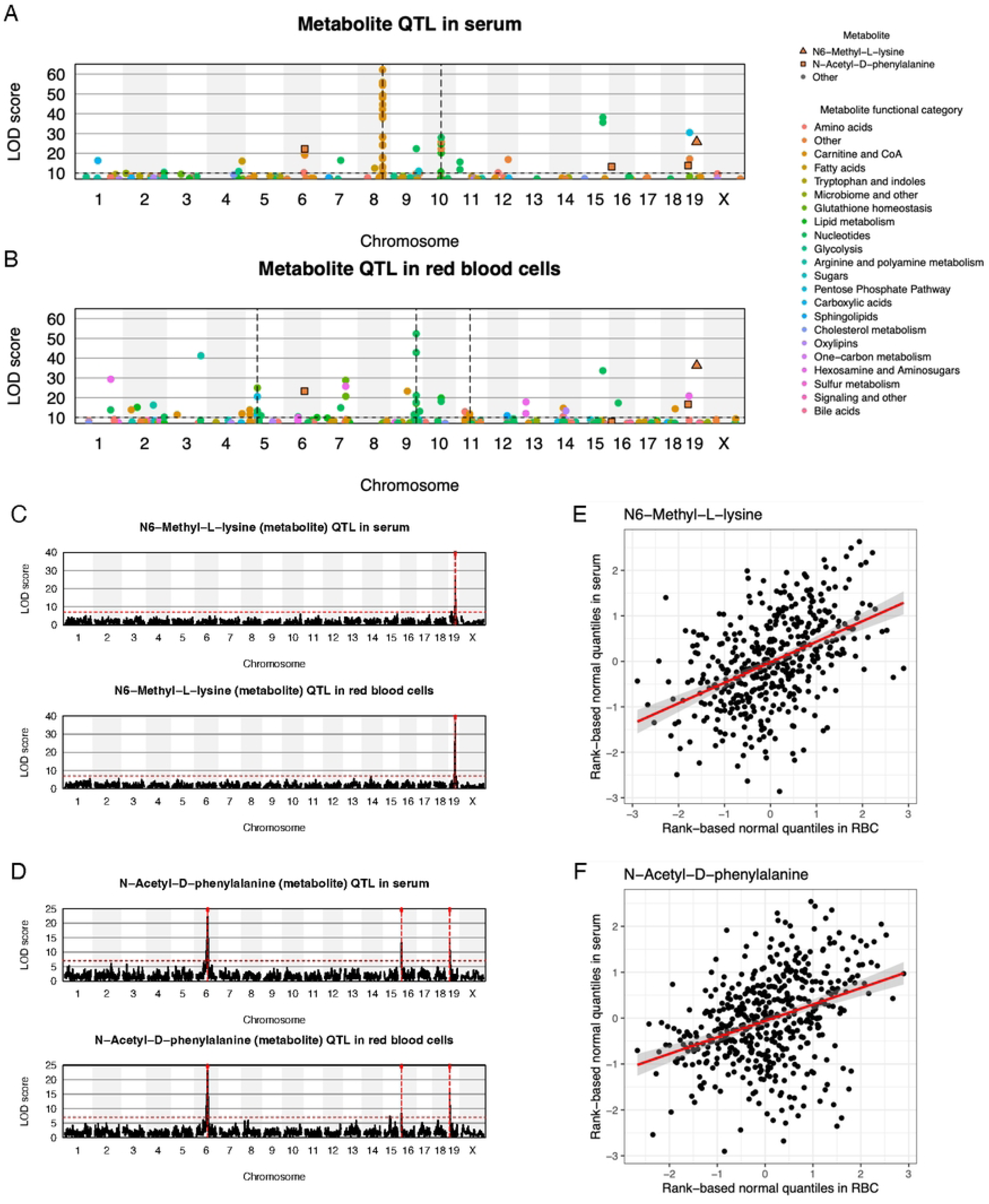
Comparison of metabolite QTL in serum and red blood cells (RBCs) reveals mostly distinct QTL networks. Manhattan plots of detected metabolite QTL (LOD score > 7) in (A) serum and RBCs. Vertical dashed lines represent QTL hotspots that are distinct for serum and RBCs. Horizontal dashed line represents a LOD score threshold of 10, used to define QTL hotspots. Four overlapping QTL between serum and RBCs are highlighted. Genome scans for (C) N6-Methyl-L-lysine and (D) N-Acetyl-D-phenylalanine, each with serum (top) and RBCs (bottom). Red vertical dashed lines represent peak QTL positions. Red horizontal dashed line represents the lenient LOD score threshold of 7. Scatter plots of (E) N6-Methyl-L-lysine and (F) N-Acetyl-D-phenylalanine, comparing metabolite levels in serum to levels in RBCs. Red best fit lines included with 95% confidence bands.

Of the 157 metabolites quantified in both serum and RBCs, only two with their 4 QTL overlap. N6-methyl-L-lysine and N-acetyl-D-phenylalanine exhibited identical QTL locations in serum and RBCs (**Figure 7C,D**), and as expected, the serum and RBC levels are correlated (**Figure 7E,F**). These findings suggest stable genetic regulation of a small subset of metabolites regardless of compartmentalization, whereas many others showed cell-type-specific QTL. This underscores the importance of tissue and cellular context in interpreting metabolic genetic regulation. We also compared sex differences and heritability, which did show some concordance between serum and RBCs (**Figure S7**),

Collectively, our high-resolution mapping in the DO mouse population identified multiple genetic loci that coordinate serum metabolite, lipid, and oxylipin profiles. These results advance our understanding of the genetic basis for circulating metabolic diversity and offer a framework for future mechanistic investigations into the roles of candidate genes in metabolic homeostasis.

## DISCUSSION

In this study, we leveraged the genetic diversity of the Diversity Outbred (DO) mouse population to map genetic determinants of serum metabolites and lipids. Using an integrated approach combining targeted and untargeted metabolomics, lipidomics, and oxylipin profiling, we identified numerous quantitative trait loci (QTL) that modulate circulating biochemical profiles. Several prominent QTL hotspots – most notably on chromosomes 8, 9, 10, and 17 – revealed strong candidate genes and gene clusters including the *Ces1*/*Ces2* carboxyl esterases and the major histocompatibility complex (MHC).

These findings extend previous mouse mQTL studies by offering high-resolution genetic mapping and by integrating multiple metabolite classes to reveal coordinated genetic control. For example, the chromosome 8 hotspots containing the *Ces1* and *Ces2* clusters were collectively associated with lipid, carnitine, and acyl-CoA species. The *Ces* cluster loci represent distinct genetic regulation, further emphasized by the distinct haplotype effects of the QTL. Their molecular profiles are also distinct, with the *Ces1* locus regulating lipids (and one oxylipin) and the *Ces2* locus regulating carnitines and acyl-CoAs. The oxylipin tauroursodeoxycholic acid (TUDCA) that mapped a QTL to *Ces1* exhibited distinct haplotype effects, suggesting substrate-specific functions among the *Ces1* paralogs. Although the *Ces* genes have historically been studied in the context of xenobiotic and prodrug metabolism [41, 42], more recent work implicates *Ces1d* and *Ces1g* in hepatic triglyceride hydrolysis and lipoprotein secretion [43–45], consistent with the lipidomic variation observed here.

Integrating data with local QTL data prioritizes *Ces1b*, *Ces1c*, and *Ces2h* as the drivers of the QTL hotpots on chromosome 8. Many of the *Ces1* family are highly expressed in the liver, though *Ces1b* was distinctly lowly expressed or absent in B6 mice. [46] Interestingly, this is consistent with the low B6 haplotype effect observed for the liver *Ces1b* local eQTL and pQTL in the DO. *Ces1c* is predominantly expressed in the liver, supporting the relevance of its liver local eQTL and pQTL data. Finally, expression of the *Ces2* family is less specific to the liver, though not totally absent, and more commonly associated with the intestine [46, 47]. However, *Ces2h* is unique and lowly expressed in the liver and distinctly highly expressed in the kidney, which is reflected in the local eQTL and pQTL data. Thus, we see convergence of evidence supporting these gene prioritizations based on matching haplotype effects between serum QTL and eQTL or pQTL along with the tissue relevance of the eQTL or pQTL.

For the chromosome 9 hotspot, *Gk5* stands out as an intriguing candidate given its known function in lipid metabolism, and thus potentially represents a novel regulator of systemic lipid homeostasis. *Gk5* encodes glycerol kinase 5, one of several paralogs that display tissue-specific expression and diverse metabolic roles. Unlike the well-characterized glycerol kinase *Gk*, *Gk5* has been studied primarily in the context of skin barrier biology and keratinocyte lipid synthesis [39]. A potential association with circulating triglycerides and phosphatidylcholines would suggest previously unrecognized roles in glycerolipid biosynthesis or turnover, possibly through tissue-specific expression in adipose or skin-derived lipoproteins. There is also some support based on a co-mapping *Gk5* local pQTL in kidney, though other genes in the QTL region are also supported.

The chromosome 10 hotspot was enriched for amino acid metabolites, particularly derivatives of tyrosine, tryptophan, and methionine. Candidate genes in this region - such as Gamma-Glutamyltransferases (*Ggt1* and *Ggt5*) and Glutathione S-Transferases (*Gstt3*, *Gstt1*, *Gstt4*, and *Gstt2*) - are involved in glutathione metabolism and processing nitrogenous compounds. The predominance of N-acetylated and oxidative catabolites suggests mitochondrial regulation of detoxification and nitrogen balance. Similar to the chromosome 9 hotspot, the local eQTL and pQTL results are complex, providing some support for Ggt5, Susd2, Gstt3, and Gstt1

Our integration eQTL and pQTL has also highlighted some of the known challenges of TWAS, such as spurious findings due to the presence of tissue data that is not trait-relevant. Alternatively, trait-relevant tissue data may not be available, and thus key local QTL not observed. Far fewer tissues are publicly available in DO mice compared to human data resources, such as GTEx [48], thus decreasing the likelihood that relevant eQTL or pQTL will be available. The QTL hotspots on chromosome 8 localize strikingly around the *Ces1* and *Ces2* gene clusters, providing strong context for relevant tissues, which ultimately allowed us to use local eQTL and pQTL to distinguish and prioritize specific carboxylesterase genes. In contrast, the genes under the hotspots on chromosomes 9 and 10 are more biologically varied in function. For example, though *Gk5* is involved in lipid metabolism and thus functionally relevant to a lipid QTL hotspot, we lack the highly relevant gene expression data for *Gk5*, notably skin tissue. It is unclear how meaningful a *Gk5* local pQTL in kidney is to lipid QTL.

Strong eQTL for gene models (e.g., *Gm16794*), which often represent pseudogenes, are likely under less selective constraint than metabolic QTL, [49] resulting in more extreme genetic effects. These extreme effects can then induce strong false positive signals in TWAS due to LD. This possibility seems likely for the chromosome 9 hotspot which exhibited LD spanning the QTL region, with multiple local QTL possessing haplotype effects with allelic series that group NZO, CAST, and PWK as a functional allele. It is important to emphasize that causal inference approaches in the context of genomics data (e.g., mediation of QTL effects [50], Mendelian Randomization [51], TWAS [52]) are always dependent on assumptions and sensitive to missing information, ultimately requiring experimental validation for final validation.

The chromosome 17 locus at 36 Mbp was the largest QTL hotspot and notably contained the MHC locus. This MHC is well known for its immunogenetic complexity and its critical role in antigen processing and presentation [53]. The locus mapped QTL exclusively for unidentified MS/MS features, the vast majority of which were consistent with representing small peptides, raising the possibility that variation in MHC genes affects systemic peptide turnover or immune surveillance. The second QTL hotspot on chromosome 17 at 58 Mbp was centered on *C3*, a key component of the complement cascade. *C3* harbors two missense variants (rs107650602 and rs52547087) with strain distribution patterns that strongly match the observed haplotype effects. Together, these two hotspots highlight extremes of how coding variation within peptides can manifest as QTL. With the *C3* locus, the QTL likely represent simple binary flags of individual coding variants. Strongly contrasting this, the MHC locus is highly polymorphic across the eight founder haplotypes, which is matched by multi-allelic QTL. This deviation from simpler bi-allelic effects pattern suggests the MHC QTL likely reflect genetic regulation of quantitative MHC-encoded peptide levels rather than the presence or absence of coding variant alleles.

The MS/MS identification analysis further supports the interpretation that a subset of untargeted features represent peptide/protein-derived circulating species. Mascot/SEQUEST searching identified 70 high-confidence peptide-spectrum matches, corresponding to 30 peptide entries and 28 unique mouse protein accessions. These identifications do not resolve all untargeted MS/MS QTL features, and we therefore retain cautious language, particularly for the MHC locus where genetic polymorphism and reference-genome limitations complicate peptide assignment. Nevertheless, the combination of peptide-like ionization states, MS/MS identifications, founder haplotype-effect patterns, and matching missense-variant strain distributions at loci such as *Apoa2* and *C3* provides a coherent genetic and biochemical framework for interpreting these signals.

Our study also builds on prior human and mouse mQTL studies by incorporating matched serum and red blood cells (RBCs) profiling from the same animals. This dual-matrix design offers the potential to reveal compartment-specific and systemic regulation of metabolite levels. Cross-matrix comparisons between serum and previously published red blood cell mQTL [20] revealed both conserved and matrix-specific genetic control. Notably, QTL for N6-methyl-L-lysine and N-acetyl-D-phenylalanine were identical across matrices, implying systemic control of their biosynthesis or clearance. *Pyroxd2* is a strong candidate gene, which is replicated by N6-methyl-L-lysine mQTL reported in humans [54]. However, the vast majority of traits exhibited matrix-specific QTL, likely underscoring metabolic demands, compartmentalization, and cellular uptake or export mechanisms. These results emphasize the importance of biological context when interpreting genetic control of metabolism.

Multi-Omics Factor Analysis (MOFA) [30] provided another view of the data by integrating the different omic layers (e.g., metabolites, lipids) into latent variables, which were then aligned with the QTL hotspots. MOFA-derived latent factors aligned well with the identified QTL hotspots, validating our locus-driven interpretations and supports the idea that coordinated polygenic regulation shapes large biochemical modules.

Several limitations warrant consideration with this study. Although DO mice capture substantial genetic diversity, their environmental exposure and evolutionary history differ from humans, limiting direct translational inference. Additionally, while our comprehensive metabolomic and lipidomic profiling offers broad coverage, untargeted workflows – while relying on MS/MS fragmentation profiles and matching against public repositories (e.g., HMDB) – may warrant further testing through orthogonal workflows (e.g., proteomics), such as for the tentative identification of MHC-derived peptides. Furthermore, despite employing rigorous statistical methods to map mQTL and integrate publicly available DO eQTL and pQTL data, our analyses cannot fully validate the true causal gene from a set of candidates within a high linkage disequilibrium (LD) QTL region. Definitive gene identification will require functional genomic approaches, including gene perturbation and metabolic flux analyses.

Despite these limitations, our results provide broad insights into the polygenic architecture of metabolic regulation. By providing an environmentally controlled reference map, this work offers a framework for interpreting human mQTL signals that are otherwise confounded by exposome heterogeneity. We identify new roles for established metabolic enzymes, propose novel candidates such as *Gk5*, and highlight immune–metabolic convergence at the MHC locus. These findings support future mechanistic work and may inform therapeutic strategies for metabolic and inflammatory diseases.

Future efforts should explore how identified genetic variants, such as for the carboxyl esterase gene clusters and *Gk5*, influence metabolic pathways at the molecular level and how environmental factors—such as diet, medication, or physiological stressors—interact with genetic predispositions. Given the known genetic variation in DO mice affecting exercise responses [14], an important next step will be determining how metabolic genetic architecture changes under various perturbations and stressors. Such work will be critical for understanding gene–environment interplay and enhancing the translational relevance of metabolomics-based genetics.

## MATERIALS AND METHODS

### DO mouse cohort

600 mice from the Diversity Outbred (DO) population [17] were acquired from the Jackson Laboratory, representing the 45th and 46th generations. This population was derived from extensive crossing of eight inbred founder strains that represent genetically distinct lineages of the house mouse: A/J, C57BL/6J, 129S1/SvlmJ, NOD/ShiLtJ, NZO/HILtJ, CAST/EiJ, PWK/PhJ, WSB/EiJ. The mice were shipped in three male-only and three female-only sub-cohorts, each comprised of 100 animals. All animals were genotyped on the GigaMUGA array [55]. 541 animals had serum samples collected, representing 298 females and 243 males. Two animals did not pass genotype quality control checks and thus removed, leaving data from 539 animals for genetic analysis. Serum samples underwent mass spectrometry-based metabolomics and lipidomics. All experimental protocols were approved by the University of Virginia IACUC on 04/22/2019 (protocol n: 4269). Mass spectrometry for metabolites, oxylipins, and lipids in red blood cells (RBCs) was described previously for this DO cohort [20, 22, 23].

### High-throughput metabolomics

Metabolomics extraction and analyses in 96 well-plate format were performed as described [56, 57]. Serum samples were overnight shipped in dry ice from the University of Virginia and transferred on ice on 96 well plate and frozen at -80 °C upon arrival at the University of Colorado Anschutz Medical Campus. Plates were thawed on ice then a 10 uL aliquot was transferred with a multi-channel pipettor to 96-well extraction plates. A volume of 90 uL of ice cold 5:3:2 MeOH:MeCN:water (*v/v/v*) was added to each well, with an electronically-assisted cycle of sample mixing repeated three times. Extracts were transferred to 0.2 µm filter plates (Biotage) and insoluble material was removed under positive pressure using nitrogen applied via a 96-well plate manifold. Filtered extracts were transferred to an ultra-high-pressure liquid chromatography (UHPLC-MS — Vanquish) equipped with a plate charger. A blank containing a mix of standards detailed before [58] and a quality control sample (the same across all plates) were injected 2 or 5 times each per plate, respectively, and used to monitor instrument performance throughout the analysis. Samples were run in randomized order within each plate, and median intra-plate normalization was performed as previously described. [20, 59] Metabolites were resolved on a Phenomenex Kinetex C18 column (2.1 x 30 mm, 1.7 um) at 45 °C using a 1-minute ballistic gradient method in positive and negative ion modes (separate runs) over the scan range 65-975 m/z exactly as previously described [56]. The UHPLC was coupled online to a Q Exactive mass spectrometer (Thermo Fisher). The Q Exactive MS was operated in negative ion mode, scanning in Full MS mode (2 μscans) from 90 to 900 m/z at 70,000 resolution, with 4 kV spray voltage, 45 sheath gas, 15 auxiliary gas. Following data acquisition, .raw files were converted to .mzXML using RawConverter then metabolites assigned and peaks integrated using ElMaven (Elucidata) in conjunction with an in-house standard library [60]. Untargeted metabolomics analyses were performed by re-running the whole sample set in MS/MS.

### Untargeted MS/MS peptide and protein identification

To evaluate whether untargeted MS/MS features mapping to QTL hotspots included peptide- or protein-derived species, MS/MS spectra corresponding to untargeted features were searched in Proteome Discoverer 3.2 using SEQUEST HT against the UniProt *Mus musculus* protein database with a target-decoy strategy. Searches allowed up to one missed cleavage, with precursor and fragment mass tolerances set to 10 ppm and 0.02 Da, respectively. No fixed modifications were specified, reflecting the absence of reduction and alkylation steps in the peptidome workflow. Variable modifications included oxidation, acetylation, deamidation, and methylation, including oxidation of Met/Cys, N-terminal and residue-level acetylation, deamidation of Asn/Gln-containing peptides, and Lys/Arg methylation. Peptide-spectrum matches were validated using Percolator, and only high-confidence assignments passing a 1% PSM-level false discovery rate were retained. This search identified 70 high-confidence peptide-spectrum matches, corresponding to 30 peptide entries mapping to 28 unique mouse protein accessions. These identifications were used to support interpretation of peptide- and protein-derived untargeted MS/MS features, but were not used to assign definitive identities to all QTL-associated MS/MS features. Features lacking confident peptide/protein matches were retained as untargeted MS/MS features and interpreted cautiously.

### High-throughput lipidomics

For lipidomics analyses, extraction procedures were identical to those described for metabolomics, though pure methanol was used. Lipid extracts were analyzed (10 µL per injection) on a Thermo Vanquish UHPLC/Q Exactive MS system using a previously described45 5 min lipidomics gradient and a Kinetex C18 column (30 x 2.1 mm, 1.7 µm, Phenomenex) held at 50 °C. Mobile phase A: 25:75 MeCN:water with 5 mM ammonium acetate; Mobile phase B: 90:10 isopropanol:MeCN with 5 mM ammonium acetate. The gradient and flow rate were as follows: 0.3 mL/min of 10% B at 0 min, 0.3 mL/min of 95% B at 3 min, 0.3 mL/min of 95% B at 4.2 min, 0.45 mL/min 10% B at 4.3 min, 0.4 mL/min of 10% B at 4.9 min, and 0.3 mL/min of 10% B at 5 min. Samples were run in positive and negative ion modes (both ESI, separate runs) at 125 to 1500 m/z and 70,000 resolution, 4 kV spray voltage, 45 sheath gas, 25 auxiliary gas. The MS was run in data-dependent acquisition mode (ddMS2) with top10 fragmentation. Raw MS data files were searched using LipidSearch v 5.0 (Thermo).

Oxylipins were extracted via a modified protein crash from previously described [61]. Starting from the same 96 well plates described above, extraction of oxylipins from serum were as follows: 10 µL of serum was aliquoted directly into 90 μL of MeOH:MeCN:H_2_O (5:3:2, v:v:v) and pipetted up and down 5x per sample. Samples were then vortexed at 4 °C for 30 minutes. Following vortexing, samples were centrifuged at 12,700 RPM for 10 minutes at 4 °C and 80 μL of supernatant was transferred to a new tube for analysis. 10 μL of extract from each sample was also combined to create a technical mixture, injected throughout the run for quality control. Lipidomics were extracted using identical preparation techniques as oxylipins, save for utilizing MeOH:IPA (1:1, v:v) instead of MeOH:MeCN:H_2_O (5:3:2, v:v:v) as an extraction solution. Mass spectrometry-based analyses were performed as previously published via a modified gradient optimized for the high-throughput analysis of oxylipins [62]. Briefly, the analytical platform employs a Vanquish UHPLC system (Thermo Fisher Scientific, San Jose, CA, USA) coupled online to a Q Exactive mass spectrometer (Thermo Fisher Scientific, San Jose, CA, USA). Lipid extracts were resolved over an ACQUITY UPLC BEH C18 column (2.1 x 100 mm, 1.7 µm particle size (Waters, MA, USA) using mobile phase (A) of 20:80:0.02 ACN:H_2_O:FA and a mobile phase (B) 20:80:0.02 ACN:IPA:FA. For negative mode analysis the chromatographic the gradient was as follows: 0.35 mL/min flowrate for the entire run, 0% B at 0 min, 0% B at 0.5 min, 25%B at 1 min, 40%B at 2.5min, 55% B at 2.6min, 70% B at 4.5 min, 100% B at 4.6 min, 100% B at 6 min, 0% B at 6.1 min and 0% B at 7 min. The Q Exactive mass spectrometer (Thermo Fisher) was operated in negative ion mode, scanning in Full MS mode (2 μscans) from 150 to 1500 m/z at 70,000 resolution, with 4 kV spray voltage, 45 sheath gas, 15 auxiliary gas. Calibration was performed prior to analysis using the PierceTM Positive and Negative Ion Calibration Solutions (Thermo Fisher Scientific).

### Technical sources of variation

Principal component analysis (PCA) was used to better understand drivers of large-scale variation in each data type (metabolites, lipids, oxylipins, and untargeted MS/MS). Briefly, outcome (e.g., metabolites, lipids, oxylipins, and untargeted MS/MS) values of zero were converted to a missing value and only outcomes with 100 non-missing observations or more were included for further analyses. Outcome matrices were log_10_ transformed and PCA performed via the pcaMethods R package [63]. PCA revealed that the first principal component (PC1) from the analyses of metabolites, oxylipins, and untargeted MS/MS reflected two distinct clusters (458 mice in cluster 1 and 83 mice in cluster 2), which were highly consistent across those data types. PC1 cluster identity was included as a covariate in all subsequent analysis for metabolites, oxylipins, and untargeted MS/MS.

### Sex differences

Sex differences in metabolites, lipids, oxylipins, and untargeted MS/MS were tested using a linear mixed effect model (LMM) that accounted for overall shared additive genetic effects [36]. Each outcome was transformed to normal quantiles for statistical analyses to reduce the influence of outlying values. For each outcome, we used the R/qtl2 package [64] to fit the following LMM:

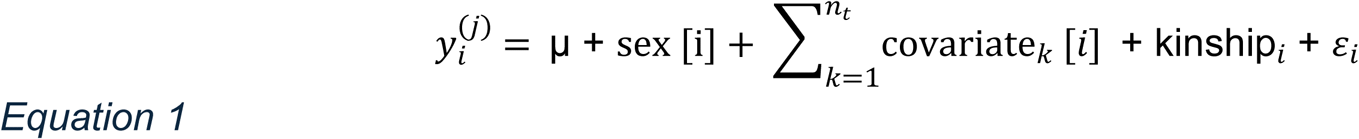

where 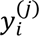 is the normal quantile for trait *j* for individual *i*, *μ* is the intercept, sex [*i*] is the contribution of sex for individual *i*, 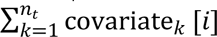 represents the summation of effects for the covariate set specific to data type *k*, kinship*_i_* and *ε_i_* are random terms representing a polygenic effect that can account for population structure and unstructured error, respectively: kinship*_i_* ∼ N(**0**, **G***τ*^2^) and *ε_i_* ∼ N(0, *σ*^2^). **G** is a *n* × *n* realized genetic relationship matrix estimated from all markers. For metabolites, oxylipins, and untargeted MS/MS, covariates included sample plate (six groups ranging from 71 to 94 mice) and PC1 cluster. For lipids, PC1 cluster was excluded. P-values were derived from a likelihood ratio test comparing the model in Equation 1 to a null model excluding sex, based on maximum likelihood parameter estimates. P-values were false discovery rate (FDR)-adjusted (P_adj_) using the Benjamini-Hochberg (BH) method [65]. Significant sex differences were defined based on a threshold of P_adj_ < 0.01. As mentioned above, sex is partially confounded with batch because animals were shipped in six separate sex-specific sub-cohorts. We adjust for batch and use a stringent significance threshold but note that this confounding could still induce false positive sex differences.

### Heritability

Narrow sense heritability (ℎ^2^), the proportion of variation explained by additive genetic effects, was estimated based on the model in Equation 1, now based on restricted maximum likelihood (REML) [66] parameter estimates. Narrow sense heritability was estimated as 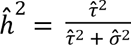 using the R/qtl2 package [64].

### QTL analysis

The QTL workflow in JAX DO mice followed previously defined conventions using the R/qtl2 package [64]. Specifically, for each outcome we performed a QTL genome scan, in which the following LMM is fit at loci spanning the genome:

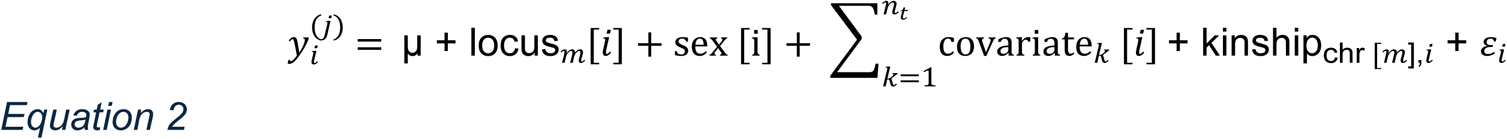

where locus*_m_*[*i*] is the effects of genetic locus *m* on individual *i*, kinship_chr [*m*],_*_i_* is a random term representing a polygenic effect that can account for population structure: kinship_chr [*m*],_*_i_* ∼ N(**0**, **G***_m_τ*^2^). **G***_m_* is an *n* × *n* genetic relationship matrix estimated from all markers excluding those from the chromosome of locus *m*, commonly referred to as leave-one-chromosome-out (LOCO) [67, 68]. LOCO increases QTL mapping power by restricting the random kinship term from absorbing signal from the tested locus term [69]. All other terms are as previously defined for Equation 1. Sex is included as a covariate now, with the locus term being the focus of statistical testing. Each genome scan was performed across a scaffold of 123,882 loci.

The conventional QTL mapping approach in multi-parental populations [15], such as the DO, is to reconstruct the genotypes at a locus in terms of the founder haplotypes rather than a SNP (eight founder haplotypes versus two SNP alleles) using a hidden Markov model and the eight founder strains’ genotypes [64, 70]. The locus term is modeled as 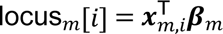 where 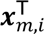 is 1 × 8 additive founder haplotype probability vector at locus *m* for individual *i* and ***β****_m_* is the 8 × 1 founder haplotype effect vector at locus *m*. During genome scans, the locus terms are estimated as fixed effects for the sake of computational efficiency. However, when interpreting founder haplotype effects at QTL peaks, the locus term is re-fit as a random effect and then estimated as best linear unbiased estimates (BLUPs) of ***β****_m_*. This conservatively constrains potentially unstable fixed effect regression coefficients.

At QTL of interest, genotypes of genetic variant (e.g., SNPs) were imputed from the founder haplotype reconstructions and the founder strain genotypes. Genetic testing is still performed based on Equation 2, but now with the locus term adjusted: locus*_m_*[*i*] = *β_m_x_i_*, where *β_m_* is the scalar effect of the allele at variant *m* and *x_m_*_,*i*_ is the genotype allele dosage at variant *m* for individual *i*. All QTL mapping, genetic variation and gene annotations were on GRCm39.

During a QTL genome scan, each locus is tested by comparing the model in Equation 2 to one excluding the locus term, which is summarized as a logarithm of odds (LOD) score statistic, analogous to a likelihood ratio. Estimating empirical significance thresholds and corresponding error rates via permutations would be computationally expensive in the context of large-scale molecular assays because the number of outcomes is large (>5,000 for this study). To avoid this and reduce the complexity of having outcome-specific significance thresholds, a lenient LOD score threshold of 7 (genome-wide type I error of 0.36 based on simulations [71]) was used to potentially capture subtle QTL, which can be supported by further biological evidence (e.g., relevant candidate genes within QTL region). For reference, a LOD score threshold of 8.3 corresponded to a genome-wide type I error of 0.05 based on simulations in this DO cohort [72]. For defining rigorous QTL hotspots, the primary focus of this study, a strict LOD score threshold of 10 was used. A QTL hotspot was defined as eight or more of such rigorous QTL mapping within 4 Mbp of each other.

### Multi-omics factor analysis (MOFA)

We performed MOFA using the MOFA2 R package [30]. We first performed a Combat [73] normalization paired with an iterative pca-based imputation (though pcaMethods R package47) to pre-adjust for covariates specific to each data layer (metabolites, lipids, oxylipins, and untargeted MS/MS). MOFA was then used with data filtered to the traits that mapped QTL to the hotspots to estimate up to 20 factors. Only factors that explain ≥1% of the variation in the data were retained. For each factor, the proportion of variance explained by each hotspot locus was estimated.

### Candidate gene prioritization via eQTL and pQTL integration

We surveyed available gene expression and protein abundance data sets from DO tissues. This resulted in 13 datasets from three broader DO cohorts (number of mice in parenthesis): liver eQTL (478) and pQTL (192) from the Svenson cohort [11]; liver eQTL (481), adipose eQTL (468), islet eQTL (378) and pQTL (378), skeletal muscle eQTL (474), and heart eQTL (483) from the Attie cohort [38, 74]; and heart eQTL (185) and pQTL (185), kidney eQTL (188) and pQTL (188), and bone eQTL (188) from the Aging cohort [75, 76]. We then integrated these data with the QTL hotspots on chromosomes 8 (*Ces1* and *Ces2* gene cluster regions), 9, and 10. For a given QTL hotspot, we took the metabolite/lipid/oxylipin with the strongest QTL (focal QTL) and used its genome scan to define a broader QTL region using a 1.5 LOD drop support interval [70]. For each gene expression and protein abundance dataset, we filtered the data to the genes encoded within the QTL support interval, performed genome scans, and detected local eQTL and pQTL using a LOD score threshold of 8 and requiring the lead marker to be within 2 Mbp of the focal QTL’s lead marker. For each local eQTL and pQTL, we estimated the haplotype effects as BLUPs from Equation 2. We then calculated the Pearson correlation coefficient between the haplotype effects of each local eQTL and pQTL and its focal QTL and tested whether it was equal to zero. We note that this approach approximates a two-sample Mendelian Randomization [77] approach, simplified to comparisons of haplotype effect summary statistics from lead markers at co-mapping QTL.

### Software

All analysis was performed using the R statistical programming language [78].

## Data availability

All data and R code to generate reported results are available at https://doi.org/10.6084/m9.figshare.31286470. Processed data and results are available at https://churchilllab.jax.org/qtlviewer/Zimring, where users can also perform interactive web-based analysis through the QTLViewer [79].

## Acknowledgements

AD and JCZ were supported by funds by the National Heart, Lung, and Blood Institute (NHLBI) (R21HL150032, R01HL146442, R01HL149714, R01HL148151). The content is solely the responsibility of the authors and does not necessarily represent the official views of the National Institutes of Health.

## Authorship Contribution

Animal studies: AH, JCZ. Metabolomics and lipidomics: TN, DS, AD. Biostatistics and Bioinformatics: GRK, CO’C, MV, GPP, GAC. pQTL analyses: GRK, GAC. Figure preparation: GRK, AD. Writing: GRK, GAC, AD. Revisions: GK, AD. Finalization: All authors.

## SUPPLEMENTAL INFORMATION

- **Document S1. Figures S1-S7.**
- **Data S1. All sex differences and heritability results, related to Figure 1**.
- **Data S2. All QTL results, related to Figure 2**.
- **Data S3. Integration of local eQTL and pQTL at QTL hotspots, related to Figures 3–5**.
- **Data S4. Mascot and Sequest search results for protein and peptide identifications based on MS/MS data**

